# Exploring the Role of YAP1 and TAZ in Pancreatic Acinar Cells and the Therapeutic Potential of VT-104 in Pancreatic Inflammation

**DOI:** 10.1101/2023.09.18.558321

**Authors:** Kevin Lopez, Janice J Deng, Yi Xu, Francis E. Sharkey, Pei Wang, Jun Liu

## Abstract

Increasing evidences have linked the hippo pathway with the fibroinflammatory diseases. We generated a series of genetic knockout mice for targeting the key components of Hippo pathway to examine the individual effects of YAP1 and TAZ on pancreatic inflammation and evaluated the therapeutic potential of the YAP1/TAZ inhibitor VT-104. Mice with acinar-specific knockout of YAP1/TAZ did not exhibit any histological abnormalities in the pancreas. LATS1/2 deficiency induced acinar-to-ductal metaplasia, immune cell infiltration and fibroblast activation, which were rescued by the homozygous knockout YAP1, but not TAZ. Additionally, treatment with VT-104 also decreased pathological alterations induced by deletions of LATS1 and LATS2 in acinar cells. Our findings highlight the critical role of YAP1 in modulating pancreatic inflammation and demonstrate that VT-104 holds therapeutic potential to mitigate pancreatitis-associated pathological manifestations. Further exploration is necessary to unravel the underlying mechanisms and translate these insights into clinical applications.

## INTRODUCTION

The importance of the fibro-inflammatory response seen during tissue damage and repair has been abundantly appreciated in recent times (1–4). This response has been demonstrated to be a complex, well-coordinated, and intricately balanced interaction between immune cells, stromal cells, and epithelial cells within damaged tissues (1). During the healing process, each of these cells harmonize through endocrine-, paracrine-, and autocrine-signaling pathways to robustly remove dead tissues via immune cells and further create supportive scaffold-like structures by stromal cells. Removal of these dead cells via macrophage and buildup of extracellular matrix (ECM) proteins via stromal cells, such as fibroblasts, promote an environment conducive to regeneration of lost epithelial tissues (2,3). Furthermore, a delicate balance of these signals is paramount to prevent over-activation of the fibro-inflammatory response which, if left uncontrolled, may yield chronic inflammation and unnecessary buildup of fibrotic tissues. These unintended byproducts not only limit the proper regeneration of damaged tissues but may also inhibit sustained and proper functioning of healthy surrounding parenchyma (4).

Within the exocrine pancreas, acinar cells account for the majority of parenchymal cells where they act as the primary producers of important digestive enzymes (5,6). During digestion, following a meal, acinar cells secrete these digestive enzymes (ex. amylase, protease, and lipase) via ducts into duodenum where they breakdown food into absorbable forms of macronutrients such as carbohydrates, proteins, and lipids (5). Though, during acute tissue injury to the pancreas, acinar cells may rupture, spilling these digestive enzymes out into the surrounding pancreatic tissue causing auto-digestive effects on the pancreas (6). In order to contain and prevent further damage, acinar cells undergo unique metaplastic changes and acquire ductal cell-like characteristics, also known as acinar-to-ductal metaplasia (ADM) (7). Immune cell infiltration, fibroblast activation, and acinar-to-ductal metaplasia are hallmark pathological characteristics of acute pancreatitis (8). When damage is sustained over time, chronic pancreatitis may develop where significant buildup of fibrotic tissue within the pancreas occurs (9). Acute and chronic pancreatitis not only limit the efficacy of pancreatic function but have also been found to increase the risk for development of pancreatic ductal adenocarcinoma (PDAC), or pancreatic cancer (10, 11). Additionally, gene mutations in acinar cells, most commonly the KRAS gene in humans, in combination with ADM has been determined to be a key factor in development of pancreatic intraepithelial neoplasia’s (PanIN), which directly precede PDAC (12,13,14).

Recently, a direct link between recurrent acute pancreatitis (AP) and chronic pancreatitis (CP) in patients has been established and is thought to be a progression of the same disease rather than independently distinct disease types (15). In addition, retrospective analysis of patients with a single event of acute pancreatitis has shown to be at increased risk for development of recurrent episodes of acute pancreatitis (16). Furthermore, pancreatitis is the leading cause of gastrointestinal-related hospital admissions proving to be a burdensome disease on people and hospitals (17). This raises the important question of how aberrant activation of the fibro-inflammatory response in the pancreas can occur and how it may be controlled.

In our previous study, we discovered the direct role of the Hippo signaling pathway in activation of the fibro-inflammatory response within acinar cells and how uncontrolled expression of YAP1/TAZ yield development of a pancreatitis-like phenotype (18). Canonically, activation of the Hippo signaling pathway in mammals inhibits cell proliferation and promotes cell apoptosis (19). These effects are primarily controlled by the core kinases, Large tumor suppressor 1 and 2 (LATS1/2), which directly phosphorylate the downstream Hippo effectors, transcriptional co-activator Yes-associated protein 1 (YAP1) and transcriptional co-activator with PDZ-binding motif (TAZ). Phosphorylation of YAP1/TAZ inhibits their cytosol-to-nuclear translocation while promoting both cytosolic retention/sequestration and ubiquitination-mediated degradation and/or glycosylation to prevent downstream transcriptional activation. In contrast, during inactivation of Hippo signaling, LATS1/2 are inactivated and no longer phosphorylate YAP1/TAZ thus allowing for YAP1/TAZ to translocate to the nucleus and participate in activation of pro-proliferative and anti-apoptotic gene transcription via interactions with TEA domain DNA-binding family members (TEAD1-4) (20,21). Though the Hippo pathway’s direct role in the fibro-inflammatory response is undoubtedly important, many questions remain unanswered in the direct, independent roles of each YAP1 and TAZ, respectively, on development of pancreatitis-like phenotypes.

Therefore, in this study, we generated novel genetically engineered mouse models to evaluate the direct roles of each YAP1 and TAZ, respectively, in activation of the fibro-inflammatory response and induction of pancreatitis-like phenotypes via the Hippo pathway. In addition, we also evaluate YAP1 and TAZ’s individual roles, through Hippo signaling, for induction of acinar-to-ductal metaplasia. Finally, we determine the efficacy of pharmaceutical inhibition of YAP1/TAZ transcriptional activities using the novel YAP1/TAZ-TEAD-complex inhibitor, VT104 (22), to determine its ability to rescue pancreatitis-associated phenotypes induced by Hippo dysregulation.

## MATERIALS AND METHODS

### Generation of conditional knockout mice

All animal study protocols were approved by the University of Texas Health San Antonio Animal Care and Use committee. Ptf1a^Cre-ERTM^ mice (The Jackson Laboratory, Bar Harbor, ME; stock number: 019378) and R26R-EYFP mice (The Jackson Laboratory; stock number: 006148) were obtained from Hebrok Lab [41]. Lats1^fl/fl^ and Lats2^fl/fl^ mice were kindly provided by Dr. Randy L. Johnson. Yap1^fl/fl^ and Taz^fl/fl^ mice were kindly provided by Dr. Eric N. Olson. We generated (1) Ptf1a^CRE-ER^ Rosa26^LSL-YFP^ mice (P), (2) Ptf1a^CRE-ER^ Lats1^fl/fl^ Lats2^fl/fl^ Rosa26^LSL-YFP^ mice (PL), (3) Ptf1a^CRE-ER^Yap1^fl/fl^ Taz^fl/fl^ Rosa26^LSL-YFP^ mice (PTY), (4) Ptf1a^CRE-ER^ Lats1^fl/fl^ Lats2^fl/fl^ Yap1^fl/fl^ Taz^fl/fl^ Rosa26^LSL-YFP^ mice (PLTY), (5)Ptf1a^CRE-ER^ Lats1^fl/fl^ Lats2^fl/fl^ Yap1^+/fl^ Taz^+/fl^ Rosa26^LSL-YFP^ mice (PLT^Hert^Y^Hert^), (6) Ptf1a^CreER^ Lats1^fl/fl^ Lats2^fl/f^ ^l^Yap1^fl/fl^ Rosa26^LSL-YFP^ mice (PLY) and (7) Ptf1a^CreER^ Lats1^fl/fl^ Lats2^fl/fl^ TAZ^fl/fl^ Rosa26^LSL-YFP^ mice (PLT). All offspring were genotyped by PCR of genomic DNA from the toe with primers specific for the Ptf1aCre-ER, Rosa26LSL-YFP, Lats1, Lats2, Yap1, and Taz transgenes. To induce the conditional knockout, 6- to 12-week-old mice were orally administrated with 180mg/kg of Tamoxifen (TAM) (Sigma-Aldrich, St. Louis, MO, T5648-5G) for 5 days, which was dissolved in corn oil (Sigma-Aldrich, St. Louis, MO, C8267). PCR was used for validation of knockout alleles.

### VT104 administration

VT104, generously supplied by Vivace Therapeutics (San Mateo, CA 94403), was prepared as previously described (22). PL mice were orally administrated TAM to knockout Lats1&2 as described above and VT104 treatment was started 72 hours after the final TAM administration. A daily dose of VT104 at 4mg/kg was administered orally, with the control group receiving the vehicle solvent. On Day 9 following the initial VT104 administration, the mice were humanely euthanized. Pancreatic tissue sections were stained with HE to quantify the severity of inflammation.

### H&E staining, immunofluorescence, and immunohistochemistry staining

**Table 1:**

| Primary Antibody | Company | Catalog # | Dilution | Application |
| --- | --- | --- | --- | --- |
| alpha-Amylase (AMY) | Sigma Aldrich | A8273 | 1:500 | IF |
| alpha-SMA (ACTA2) | Santa Cruz Biotechnology | 53142 | 1:250 | IF |
| CD45 | Biolegend | 103102 | 1:50 | IHC |
| KRT19 | DSHB | Troma-III | 1:50 | IF |
| F4/80 | Cell Signaling Technology | 70076S | 1:50 | IF |
| GFP (YFP) | Aves | GFP-1020 | 1:500 | IF |
| SPP1 | RD Systems | AF808 | 1:50 | IF |
| TAZ | Sigma Aldrich | HPA007415-100UL | 1:50 | IF |
| YAP | Cell Signaling Technology | 14074S | 1:100 | IF |

**Table 2:**

| **Secondary Antibody** | **Company** | **Catalog #** | **Dilution** | **Application** |
| --- | --- | --- | --- | --- |
| Alexa Fluor® 488-conjugated AffiniPure Donkey Anti-Mouse IgG (H+L) | Jackson ImmunoResearch | 715-545-150 | 1:250 | IF |
| Alexa Fluor® 647-conjugated AffiniPure Donkey Anti-Mouse IgG (H+L) | Jackson ImmunoResearch | 715-605-150 | 1:250 | IF |
| Cy™3-conjugated AffiniPure Donkey Anti-Mouse IgG (H+L) | Jackson ImmunoResearch | 715-165-150 | 1:250 | IF |
| Alexa Fluor® 647-conjugated AffiniPure Donkey Anti-Rabbit IgG (H+L) | Jackson ImmunoResearch | 711-605-152 | 1:250 | IF |
| Cy™3-conjugated AffiniPure Donkey Anti-Rabbit IgG (H+L) | Jackson ImmunoResearch | 711-165-152 | 1:250 | IF |
| Alexa Fluor® 647-conjugated AffiniPure Donkey Anti-Rat IgG (H+L) | Jackson ImmunoResearch | 712-605-150 | 1:250 | IF |
| Cy™3-conjugated AffiniPure Donkey Anti-Rat IgG (H+L) | Jackson ImmunoResearch | 712-165-150 | 1:250 | IF |
| Alexa Fluor® 647-conjugated AffiniPure Donkey Anti-Goat IgG (H+L) | Jackson ImmunoResearch | 705-605-147 | 1:250 | IF |
| Cy™3-conjugated AffiniPure Donkey Anti-Goat IgG (H+L) | Jackson ImmunoResearch | 705-165-003 | 1:250 | IF |
| Alexa Fluor® 488-conjugated AffiniPure Donkey Anti-Chicken IgY (IgG) (H+L) | Jackson ImmunoResearch | 703-545-155 | 1:250 | IF |
| Peroxidase-conjugated AffiniPure Goat Anti-Rat IgG (H+L) | Jackson ImmunoResearch | 112-035-003 | 1:250 | IHC |

### Statistical analysis

All results in this study were presented as the mean ± standard error of the mean (s.e.m.). Statistical analysis was performed by a two-tailed Student’s t-test. All p-values < 0.05 were considered statistically significant.

## RESULTS

### Deletion of YAP1/TAZ in acinar cells does not induce abnormal or pancreatitis-like phenotype long-term

Previously, we evaluated the effects of homozygous YAP1 and TAZ deletions in adult mice acinar cells and determined that no abnormal phenotypes or pathologies arose from loss of YAP1/TAZ genes in two weeks after Tamoxifen (TAM) administration (18). However, whether YAP/TAZ deletion has long term effect on pancreas homeostasis was not clear. To answer this, we administered TAM daily for five consecutive days by oral gavage at the concentration of 180mg/kg to delete the floxed alleles in acinar cells of the *Ptf1a^CreER^Yap1^fl/fl^Taz^fl/fl^Rosa26^LSL-YFP^*(PTY) mice (**Fig 1A**). Then, we sacrificed the mice at 4 months post-TAM administration. Histological analysis by H&E stain revealed no abnormalities of the pancreatic tissues (n=6) **(Fig 1B)**. In conclusion, these data suggest that YAP1/TAZ are dispensable for the long-term homeostasis of acinar cells under physiological conditions.

**Figure 1:**
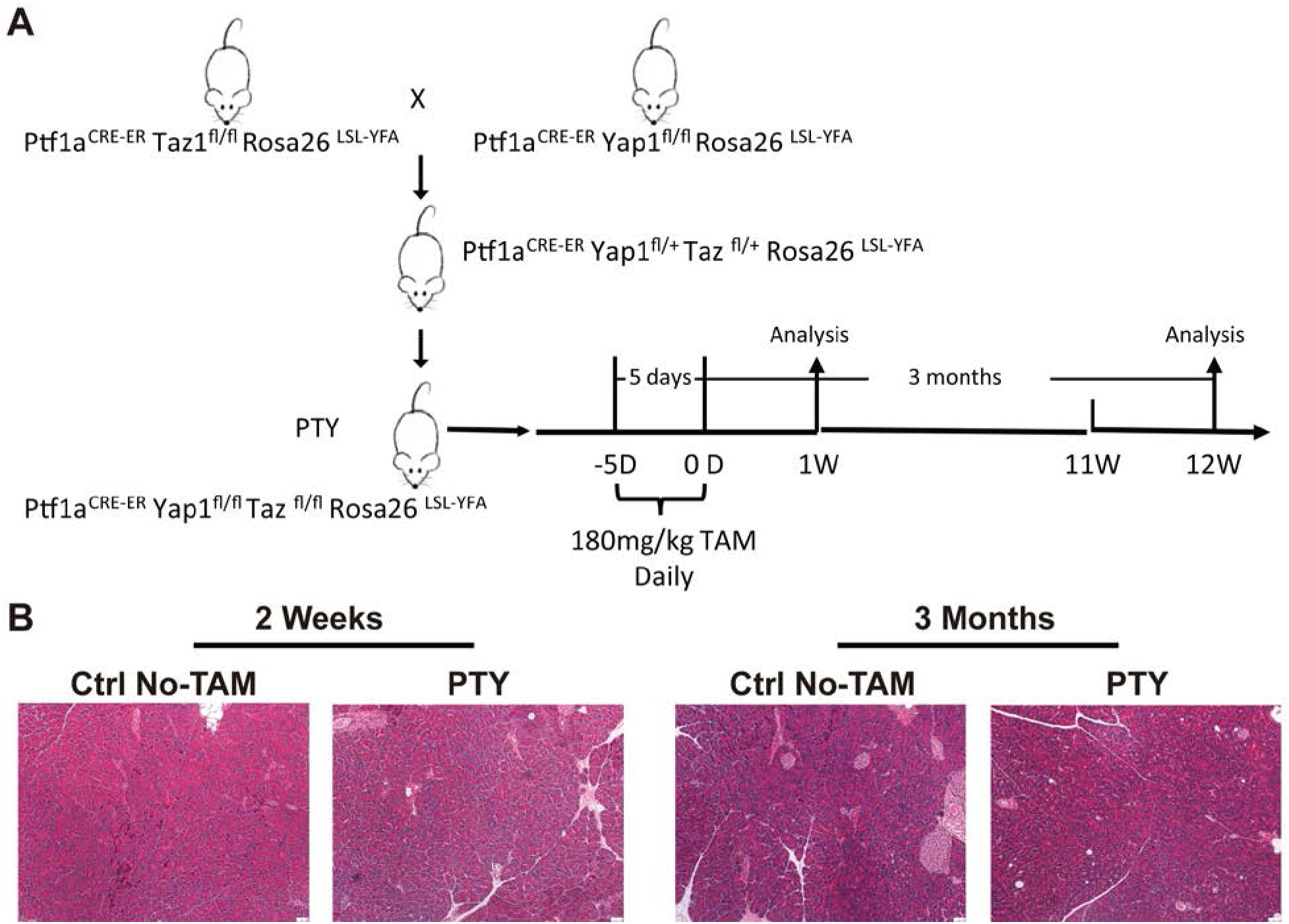
Acinar-specific knockout of YAP1 and TAZ did not disturb pancreas homeostasis under normal conditions. A) The schema of experimental procedure. B) The representative H&E staining images of the pancreas collected from the mice treated as indicated.

### Heterozygous YAP1/TAZ knockout in acinar cells does not rescue Lats1/2 knockout-induced pancreatic damage and inflammation

Our previous study found that homozygous YAP1 and TAZ knockout in Lats1/2-null acinar cells in adult mice rescued pancreatic damage and inflammation (18). In developping pancreas, heterozygous YAP1 knockout largely reversed the defects triggered by Hippo deficiency (23). To confirm the similar effect in adult pancreas, we analyzed the pancreas of heterozygous YAP1 and TAZ knockout in Lats1/2-null acinar cells (*Ptf1a^CreER^Lats1^fl/fl^Lats2^fl/fl^Yap1^fl/+^Taz^fl/+^Rosa26^LSL-YFP^*, PLThYh) on pancreatic damage and inflammation **(Supplemental Fig 1)**. As expected, we did not observe abnormality in control mice without TAM administration by H&E stain (n=3) **(Fig 2A)**. The *Ptf1a^CreER^Lats1^fl/fl^Lats2^fl/fl^Rosa26^LSL-YFP^*(PL) mice also received the TAM administration. Histological analyses at 7 days post-TAM administration by H&E stain revealed severe pancreatic damage of PLThYh mice (n=7) that was indistinguishable from the defects observed in PL mice (n=5) **(Fig 2A)**. Immunofluorescent staining demonstrated robust F4/80+ macrophage infiltration and fibroblast activation, measured by α-SMA staining, at similar levels in both PL and PLThYh mice **(Fig 2B)**. In addition, immunohistochemistry stain of marker of proliferation Ki67 also had indistinguishable differences between PL and PLThYh mice **(Fig 2C).** Further immunofluorescent staining of Amylase and cytokeratin 19 (CK19) revealed extensive, indistinguishable levels of ADM in the pancreas from both PL and PLThYh mice **(Fig 2D).** Altogether, the phenotypes observed in PL and PLThYh mice demonstrated inability of YAP1 and TAZ haplotypes to rescue pancreatic damage and inflammation induced by Hippo-pathway disruptions in adult mice acinar cells.

**Figure 2:**
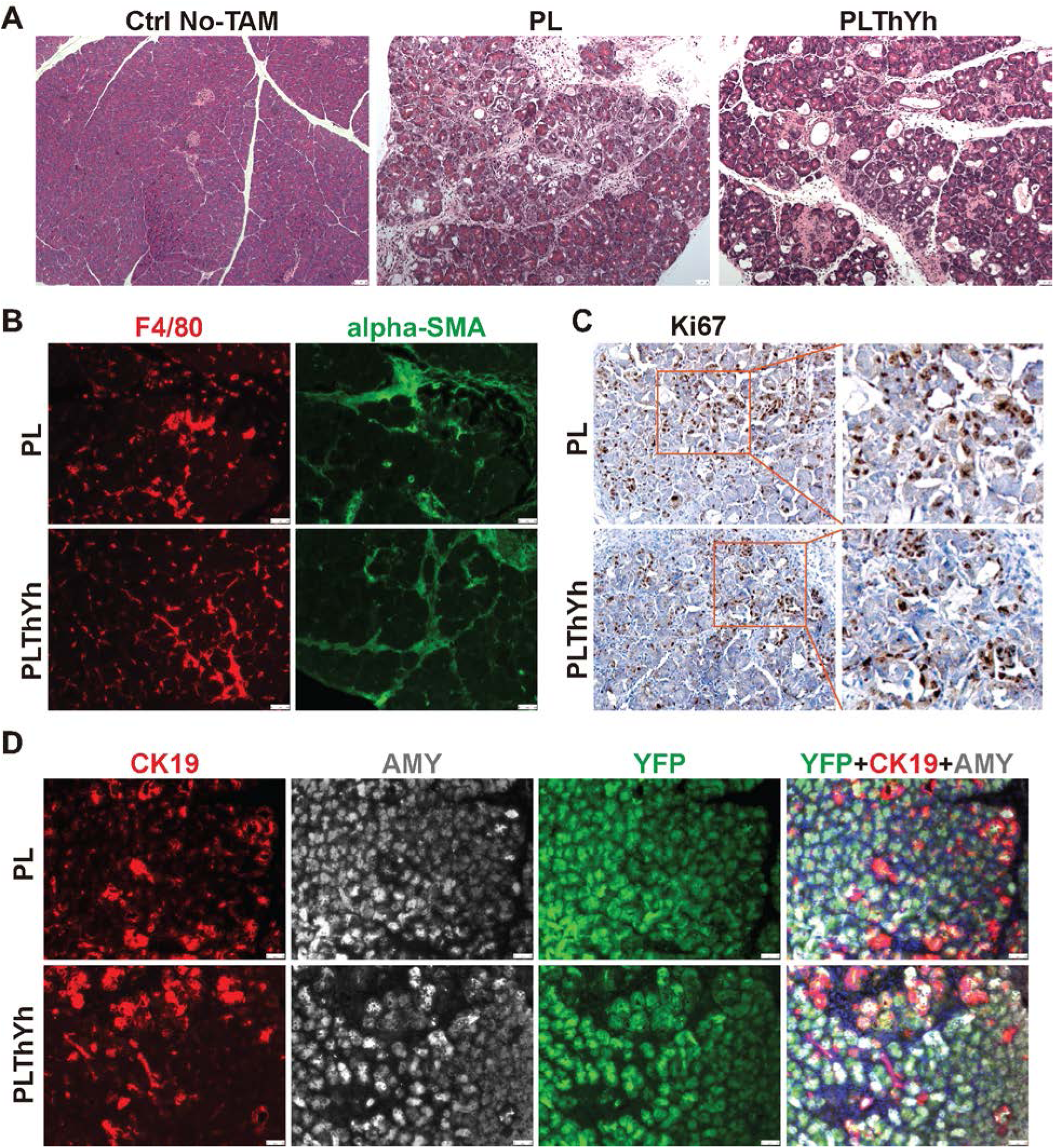
The pancreatic inflammation and damage caused by acinar-specific LATS1/2 knockout were not mitigated by the haplotype of YAP1 and TAZ. A) The representative H&E staining images of the pancreas collected from the mice treated as indicated. B) The representative IF images of F4/80 and alpha-SMA in the pancreas collected from PL and PLT^Hert^Y^Hert^ mice 7 days after final TAM administration. C) The representative IHC images of Ki67 in the pancreas collected from PL and PLY mice 7 days after final TAM administration. D) The representative IF images of Amylase and CK19 in the pancreas collected from PL and PLT^Hert^Y^Hert^ mice 7 days after final TAM administration.

### Homozygous YAP1 knockout rescues of pancreatic inflammation and acinar-to-ductal metaplasia induced by acinar cell-specific Lats1/2 deletions

Although some early studies suggested the redundant functions of YAP1 and TAZ (24), emerging evidence suggests that they may play synergistic or even opposing roles in different biological contexts (25). In order to evaluate the contributions of YAP1 activation on pancreatic inflammation induced by acinar-specific Lats1/2-knockout in adult mice pancreas, *Ptf1a^CreER^Lats1^fl/fl^Lats2^fl/fl^Yap1^fl/fl^Rosa26^LSL-YFP^*(PLY) mice were created **(Supplemental Fig 1)** and administered TAM as previously described to simultaneously delete LATS1/2 and YAP1 in acinar cells. Upon histological analysis by H&E stain at day 6 post-TAM administration, PLY mice (n=6) showed a near complete resolution of overall pancreatic damage and inflammation compared to PL mice (n=3), though containing minor sporadic lesions **(Fig 3A)**. Next, we evaluated the status of nuclear/cytosolic YAP1 and TAZ using immunohistochemistry, which was seen to be consistent with genotyping for PL and PLY mice. In PL pancreas, strong nuclear YAP1 and TAZ expression was seen in acinar cells **(Fig 3B)**. In PLY pancreas, non-damaged areas showed absence of nuclear and cytosolic YAP1, while nuclear and cytosolic expression of TAZ persisted **(Fig 3B)**. Interestingly, all small sporadic lesions seen in PLY mice showed strong nuclear YAP1 expression, suggesting incomplete knockout of YAP1 (**Supplemental Fig 2A**). In addition, PLY pancreas had reduced expression of downstream YAP1 target SPP1 compared to PL mice **(Fig 3B),** further supporting the successful deletion of YAP1. The knockout of YAP1 also significantly decreased macrophage infiltration and activation of fibroblasts revealed by immunofluorescent staining for F4/80 **(Fig 3C)**. Quantification of average positive fluorescent area of F4/80+ macrophage in PL mice (n=3) and PLY mice (n=3) showed a statistically significant decrease in macrophage recruitment. For α-SMA stain quantification, the average positive fluorescent area of α-SMA for PL and PLY mice also showed a statistically significant decrease in fibroblast activation **(Fig 3C)**. The PLY pancreas also had significantly less Ki67-positive cells compared to PL mice (**Fig 3D**). In addition, the numbers of acinar cells undergoing ADM which were evaluated by GFP, CK19, and Amylase immunofluorescent co-staining were significantly less in PLY mice than PL mice, demonstrating reduced occurrence of ADM. Quantification of average ADM using triple-positive GFP/CK19/Amylase cells showed that there were less ADM in PLY pancreas compared with PL pancreas **(Fig 3E).** We further examined the co-relation of YAP1 activation and CK19 induction in acinar cells. We clearly identified sporadic GFP+ YAP1+ cells in PLY mice, consistent with the conclusion that YAP1 has not been completely ablated in all acinar cells. We did not observe any GFP+ CK19+ ADM cells without nuclear YAP1 expression, suggesting that YAP1 activation was essential for CK19 induction in acinar cells. On the other hand, we noticed that some GFP+YAP1+ cells lacked CK19 expression **(Supplemental Fig 2B),** suggesting that additional signals might be required for YAP1 to promotes CK19 expression in Lats1/2-null acinar cells. Altogether, these data suggest that YAP1 is the essential driving factor that mediates the inductive effects of acinar-specific LATS1/2 inactivation on pancreatic damage, inflammation, and ADM.

**Figure 3.**
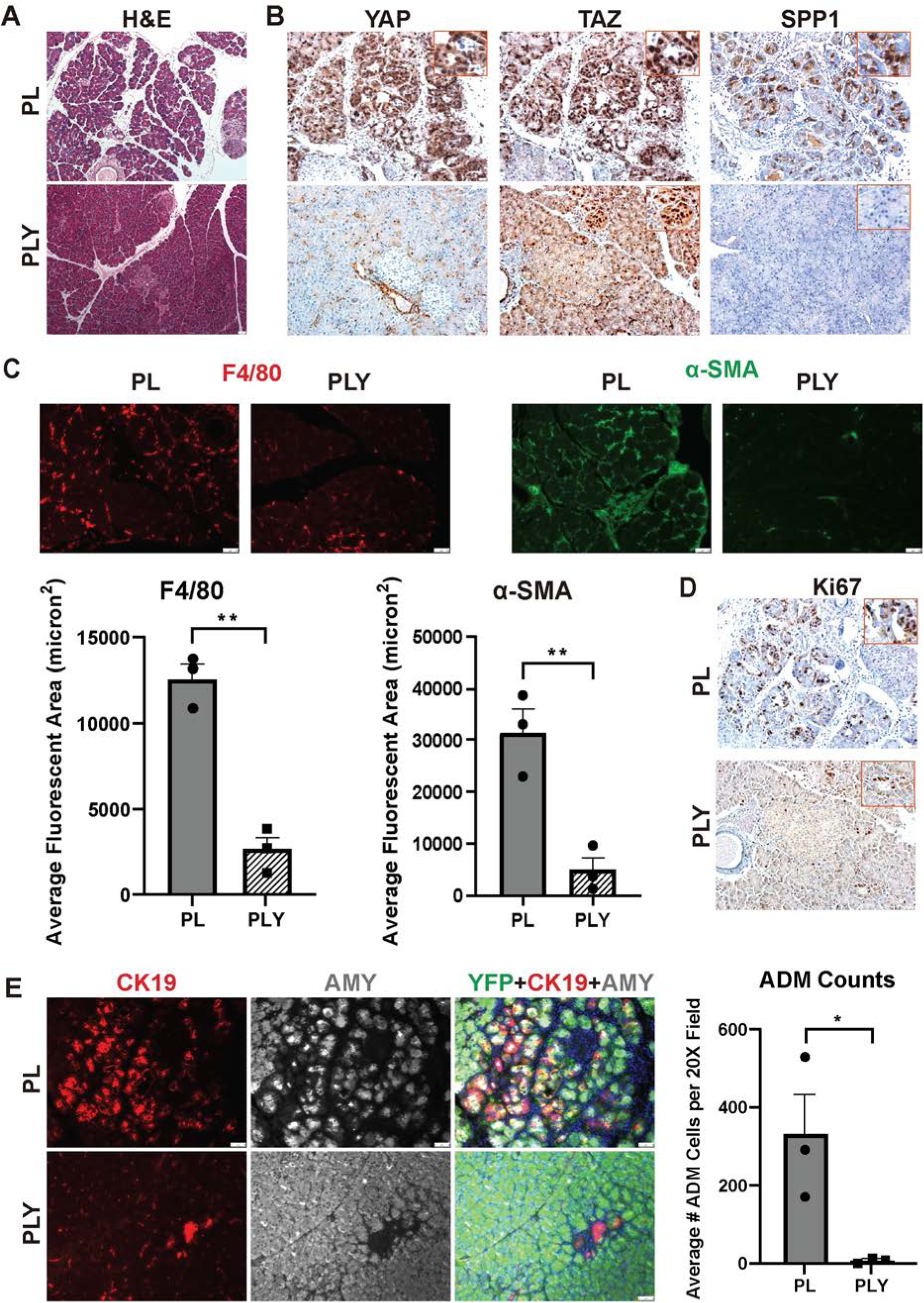
YAP1 is the essential mediator of the pancreatic inflammation and damages induced by LATS1/2 inactivation in acinar cells: A) The representative H&E staining images of the pancreas collected from PL and PLY mice 7 days after final TAM administration. B) The representative IHC images of YAP1, TAZ and SPP1 in the pancreas collected from PL and PLY mice 7 days after final TAM administration. C) The representative IF images of F4/80 and alpha-SMA in the pancreas collected from PL and PLY mice 7 days after final TAM administration. Average positive fluorescent area (in squared microns) of F4/80+ macrophage in PL mice (n=3) was 12624±1512 while PLY mice (n=3) was 2643±1330 with p-value =0.001. The average positive fluorescent area of α-SMA for PL mice was 31576±7961 and PLY mice was 4860±4315 with p-value = 0.007. D) The representative IHC images of Ki67 in the pancreas collected from PL and PLY mice 7 days after final TAM administration. E) The representative IF images of GFP, CK19, and Amylase in the pancreas collected from PL and PLY mice 7 days after final TAM administration. Quantification of average ADM numbers per 20X field using triple-positive GFP/CK19/Amylase cells in PL mice was 330±182.6 while PLY mice was 7±6.557 with p-value =0.037

### Homozygous TAZ knockout insignificantly reduced pancreatic inflammation and acinar-to-ductal metaplasia induced by acinar-specific Lats1/2 deletions

Although our data demonstrated that TAZ activation in acinar cells was insufficient to cause the pancreatic inflammation and damage induced by acinar-specific Lats1/2-knockout in adult mice pancreas, it remained unclear whether TAZ activation was required for these pathological alterations. To this end, we generated *Ptf1a^CreER^Lats1^fl/fl^Lats2^fl/fl^Taz^fl/fl^Rosa26^LSL-YFP^*(PLT) mice **(Supplemental Fig 1)** and administered TAM as previously described to knockout LATS1/2 and TAZ. Upon histological analysis at day 6 post-TAM administration, H&E stain of PLT mice (n=3) showed pronounced pancreatic damage and inflammation resembling PL mice (n=3) **(Fig 4A)**. We evaluated the status of nuclear/cytosolic TAZ and YAP1. In the pancreas of PLT mice, nuclear YAP1 expression was seen in acinar cells, like PL mice, while TAZ expression was generally absent **(Fig 4B)**. In some lesions of PLT mice, nuclear TAZ expression was observed, suggesting incomplete knockouts in these cells. Nevertheless, TAZ expression was dispensable in cells within lesions **(Fig 4B)**.In contrast, all lesions in PLT mice clearly expressed nuclear YAP, which was consistent with nuclear YAP staining in the lesions observed in PLY mice **(Fig 3B)**. Additionally, immunohistochemistry stains of SPP in PLT mice were closely resembled that of PL mice **(Fig 4B)**. Immunofluorescent stains revealed the PLT mice mimicked PL mice with increased F4/80+ macrophage infiltration and alpha-SMA+ fibroblast activation **(Fig 4C)**. The quantification of F4/80 and alpha-SMA stains yielded no statistically significant changes between PL and PLT mice. Similarly, the PLY pancreas also had similar numbers of Ki67-positive cells to PL mice **(Fig 4D**). Next, we evaluated the occurrence of ADM by evaluating GFP, CK19, and Amylase immunofluorescent co-staining as previously described. The PLT mice demonstrated similar occurrence of ADM to PL mice **(Fig 4E)**. Though average ADM numbers slightly decreased from PL to PLT mice, there was no statistically significant difference. We further examine the co-relation of AMD with YAP1 activation. In PLT mice, nuclear YAP1 staining was present in the majority of GFP-labeled acinar cells, which was consistent with the previous immunohistochemistry YAP1 stain on PLT mice **(Fig 4B)**. As expected, all GFP+ CK19+ ADM cells in the PLT mice expressed nuclear YAP1 **(Supplemental Fig 2C)**. Together, these data not only suggest that TAZ has a minimal effect on induction of pancreatic inflammation and is dispensable for ADM in Lats1/2-null acinar cells, but also further confirmed that YAP1 is the primary initiator and driver of these responses within the pancreas.

**Figure 4:**
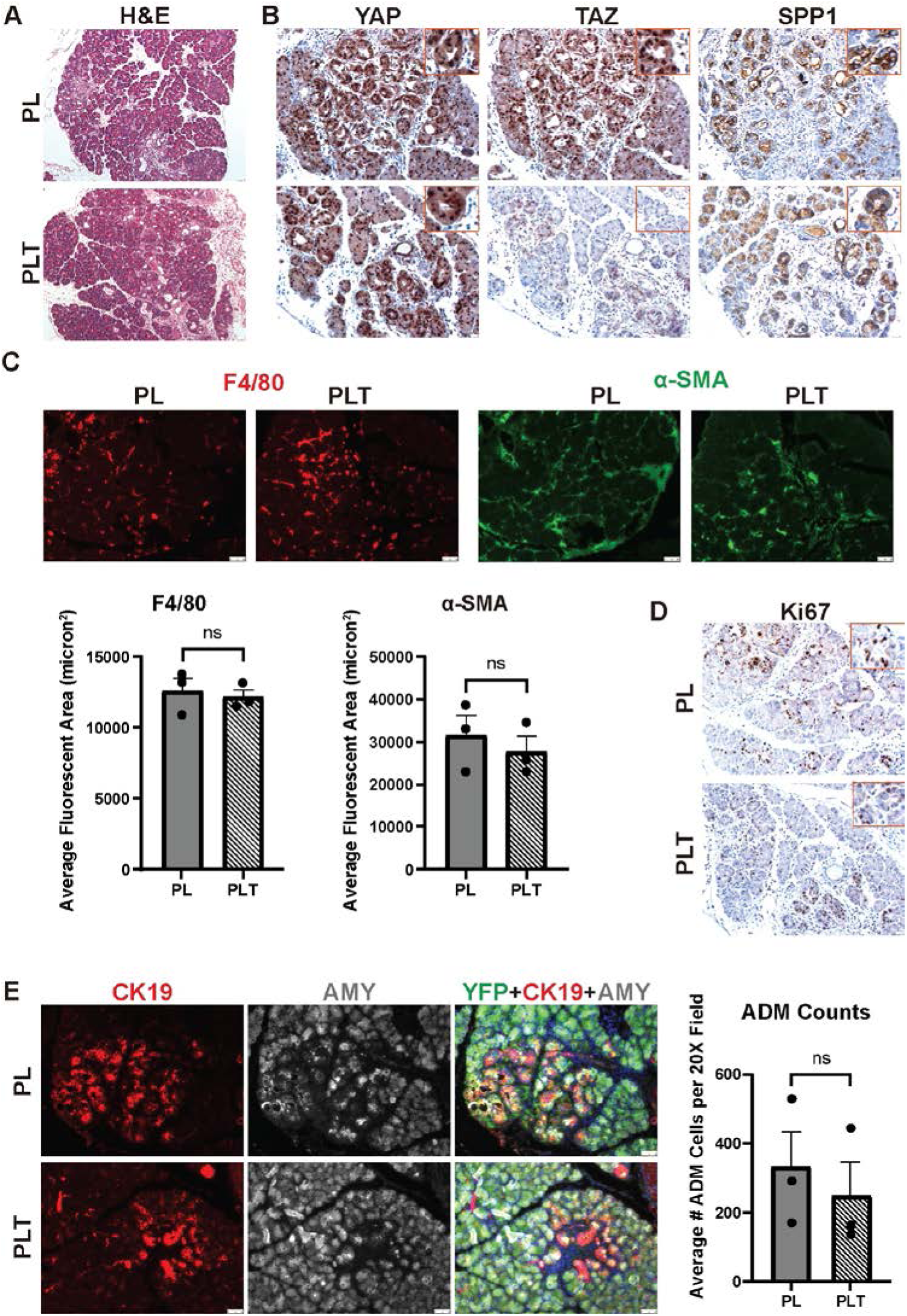
TAZ is dispensable for the induction of pancreatic inflammation and damages in mice with LATS1/2 deficiency in acinar cells. A) The representative H&E staining images of the pancreas collected from PL and PLT mice 7 days after final TAM administration. B) The representative IHC images of YAP1, TAZ and SPP1 in the pancreas collected from PL and PLT mice 7 days after final TAM administration. C) The representative IF images of F4/80 and alpha-SMA in the pancreas collected from PL and PLT mice 7 days after final TAM administration. D) The representative IHC images of Ki67 in the pancreas collected from PL and PLT mice 7 days after final TAM administration. E) The representative IF images of GFP, CK19, and Amylase in the pancreas collected from PL and PLT mice 7 days after final TAM administration.

### Pharmaceutical YAP1/TAZ-TEAD-complex inhibitor VT104 treatment attenuates inflammation and ADM induced by acinar-specific Lats1/2 deletions

Previously, Tang et al. identified novel YAP/TAZ-TEAD-complex inhibitor VT104 to be a promising and potent pharmaceutical candidate for cancer treatment due its ability to inhibit tumor growth and proliferation of NF2-deficient Mesothelioma in vitro and in xenograft in vivo models (22). We demonstrated the requirement of YAP1 to induce pancreatitis-like phenotypes through genetic mouse models. To further evaluate if a pharmacologic inhibition of YAP1-TEAD interaction can attenuate pancreatic inflammation and damage due to inactivation of Hippo pathway, we treated TAM-administrated PL mice with Saline (n=3) as controls or VT104 (n=5) at a concentration of 3mg/Kg for 9 days **(Fig 5A)**. Expectedly, histological analysis by H&E stain revealed extensive damage to PL+Saline treatment mice consistent with previous PL mice phenotypes. In contrast, the PL+VT104 treatment mice showed significant reduction in overall pancreatic damage and inflammation **(Fig 5B)**. We further performed immunofluorescent stains on F4/80+ macrophage and alpha-SMA of activated fibroblasts, each showing significant reductions after VT104 treatment compared to Saline-treated mice, respectively. Quantification of F4/80 showed the average fluorescent area (in squared microns) for PL+Saline (n=3) was 41338±5096 while PL+VT104 (n=5) was 8206±3601 with a p-value <.0001. Quantification of alpha-SMA showed the average fluorescent area for PL+Saline was 65252±10555 while PL+VT104 was 20030±10923 with a p-value = 0.001 **(Fig 5B)**. We then analyzed ADM numbers using the same parameters previously described by immunofluorescent co-stains where PL+VT104 treatment mice demonstrated significant reduction in ADM **(Fig 5C)**. Quantification of ADM numbers for PL+Saline mice were 543.3±81.45 while PL+VT104 mice were 135.2±85.27 with a p-value = 0.001 **(Fig 5C)**. Altogether, these data demonstrate the robust efficacy of pharmaceutically targeted YAP/TAZ-TEAD-complex inhibition in resolving severe pancreatitis phenotypes induced by genomic-level Hippo-pathway inactivation.

**Figure 5:**
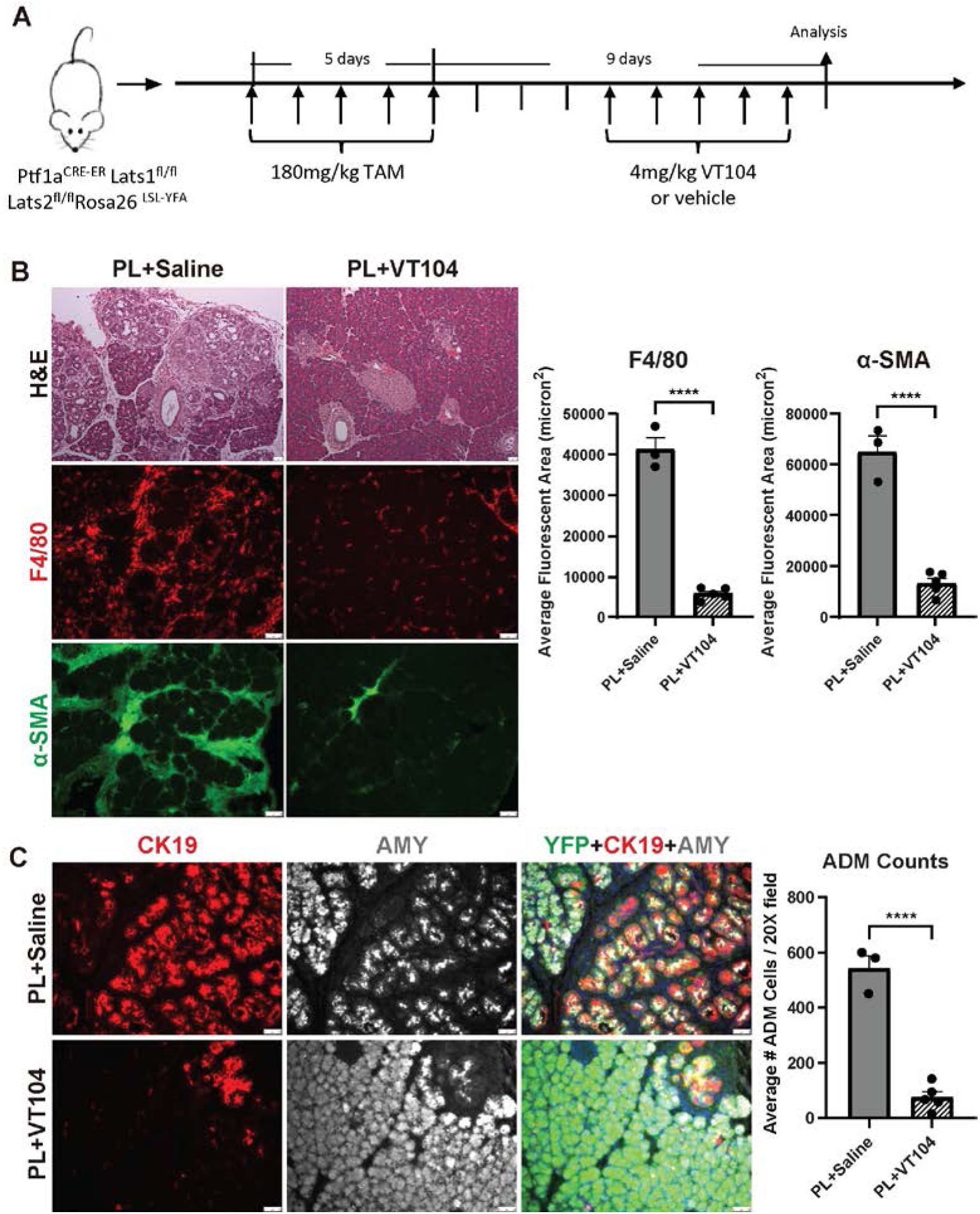
Yap1/Taz pharmaceutic inhibitor VT104 ameliorate pancreatic damages induced by Lats1/2 ablation. A) The schema of VT104 treatment procedure. B) The representative H&E staining images of the pancreas collected from PL and PLT mice 7 days after final TAM administration. C) The representative IF images of F4/80 and alpha-SMA in the pancreas collected from PL and PLT mice 7 days after final TAM administration. D) The representative IF images of GFP, CK19, and Amylase in the pancreas collected from PL and PLT mice 7 days after final TAM administration.

## DISCUSSION

We previously reported that loss Lats1&2 in mouse adult acinar cells led to inflammation in pancreas which can be rescued by further deletion of YAP1 and TAZ (18). YAP1 and TAZ were originally considered as the redundant down-stream mediators of Hippo pathway (24), but subsequent studies have challenged this notion (25). The whole-body knockout of YAP1 in mice resulted in embryonic lethal (26), whereas the knockout of TAZ was not lethal during development, although it did lead to abnormality in certain tissues (27). These observations underscore the importance of assessing the individual functions of YAP1 and TAZ in context-dependent manner. Here we showed that deletion of YAP1 but not TAZ can rescue the fibroinflammation caused by loss of Lats1/2. These observations demonstrated that YAP1 activation is necessary and sufficient in facilitating the changes induce by the deletions of LATS1/2 in adult pancreatic acinar cells. In addition, our data also exclude the possibility that TAZ activation in LATS1/2-deficient acinar cells is suppressed by YAP1 knockout, suggesting that TAZ activation was neither required nor sufficient to mediate the changes induced by LATS1/2 deletions in adult pancreatic acinar cells.

The distinct consequences of YAP1 and TAZ activation reflect their differential transcriptional activities in adult pancreatic acinar cells. It is important to note that both YAP1 and TAZ need to interact with other factors to initiate transcription, as they do not bind DNA directly. The TEADS are the most well-known DNA binding transcriptional factors (TF) that interact with YAP1 and TAZ (20,21). However, YAP1 and TAZ may use other DNA binding TFs to regulate gene expression (28). While it is possible that YAP1 has TEAD-independent targets that may contribute to the induction of fibroinflammation, the earlier studies showed CTGF, a well-known TEAD-dependent target of YAP1 and TAZ, was partially responsible for it (18, 29). Consistent with this view, treatment with VT-104, a new compound which can block the interaction between TEADs and YAP1, largely rescue the pancreatic damages in mice with acinar-specific deletions of LATS1/2.These findings highlight the therapeutic potential of VT-104 for the treatment of pancreatitis.

The observation that heterozygous YAP1 knockout had minimal effects to lessen the inflammation caused by LATS1/2 deficiencies, suggesting that the acinar cells are highly sensitive to YAP1 activation, which should be tightly controlled in physiological conditions. Indeed, in the absence of stresses, no detectable pancreatic abnormalities were observed at either 1 week or 4 months after YAP1 and TAZ knockout in pancreatic acinar cells. Therefore, it is crucial to delve deeper into the roles of YAP1 activation and investigate how it is exploited to facilitate disease progression. The roles of YAP1 in driving ADM remain controversial. One previously study suggested that YAP1 activation is sufficient to induce ADM (30). This conclusion, however, relied partially on observations from *in vitro* culture models. In our *in vivo* genetic models, we observed that some acinar cells with nuclear YAP1 did not express CK19, suggesting additional factors are also required to induce typical ADM. However, the observation that all acinar-derived CK19 positive cells displayed nuclear localization of YAP1 emphasized the important contributions of YAP1 activation to ADM phenotypes. Interestingly, previous studies have reported that in the pancreatic cancer model, oncogenic KRAS perpetuated the acute injury-induced ADM in YAP1-dependent manner (31, 32). In conjunction with the observation that oncogenic KRAS can stabilize YAP1 activation at post-translation level (31), these data suggested that YAP1 activation played the key roles in stabilizing the duct-like phenotypes in ADM lesions.

In summary, we developed multiple genetic models to systemically evaluate the roles of Yap1 and Taz in in driving the pancreatic pathological changes resulting from the disruption of the Hippo pathway within pancreatic acinar cells. Additionally, we explored the potential of mitigating these alterations through pharmaceutical inhibition of the transcriptional activity of Yap1/Taz. Therefore, aside from clarifying the distinct roles of Yap1 and Taz in acinar cells and shedding light on understanding the roles of Yap1 in ADM development, a crucial alternation in pancreatic disease progression, our research also lays the groundwork for the rationale for pharmaceutically targeting Yap1 as a promising strategy to treat pancreatic diseases.

## Author Contributions

Conceptualization: Wang P, Liu J

Data curation: Lopez K, Deng J, Xu Y

Formal analysis: Lopez K, Liu J Sharkey F, Xu Y

Funding acquisition: Wang P, Liu J

Methodology: Lopez K, Liu J Sharkey F, Xu Y

Project administration: Wang P, Liu J

Supervision: Wang P, Liu J

Writing – original draft: Lopez K, Wang P, Liu J

Writing – review & editing: Wang P, Liu J, Xu Y

## Supplementary data

**Supplementary Figure 1:**
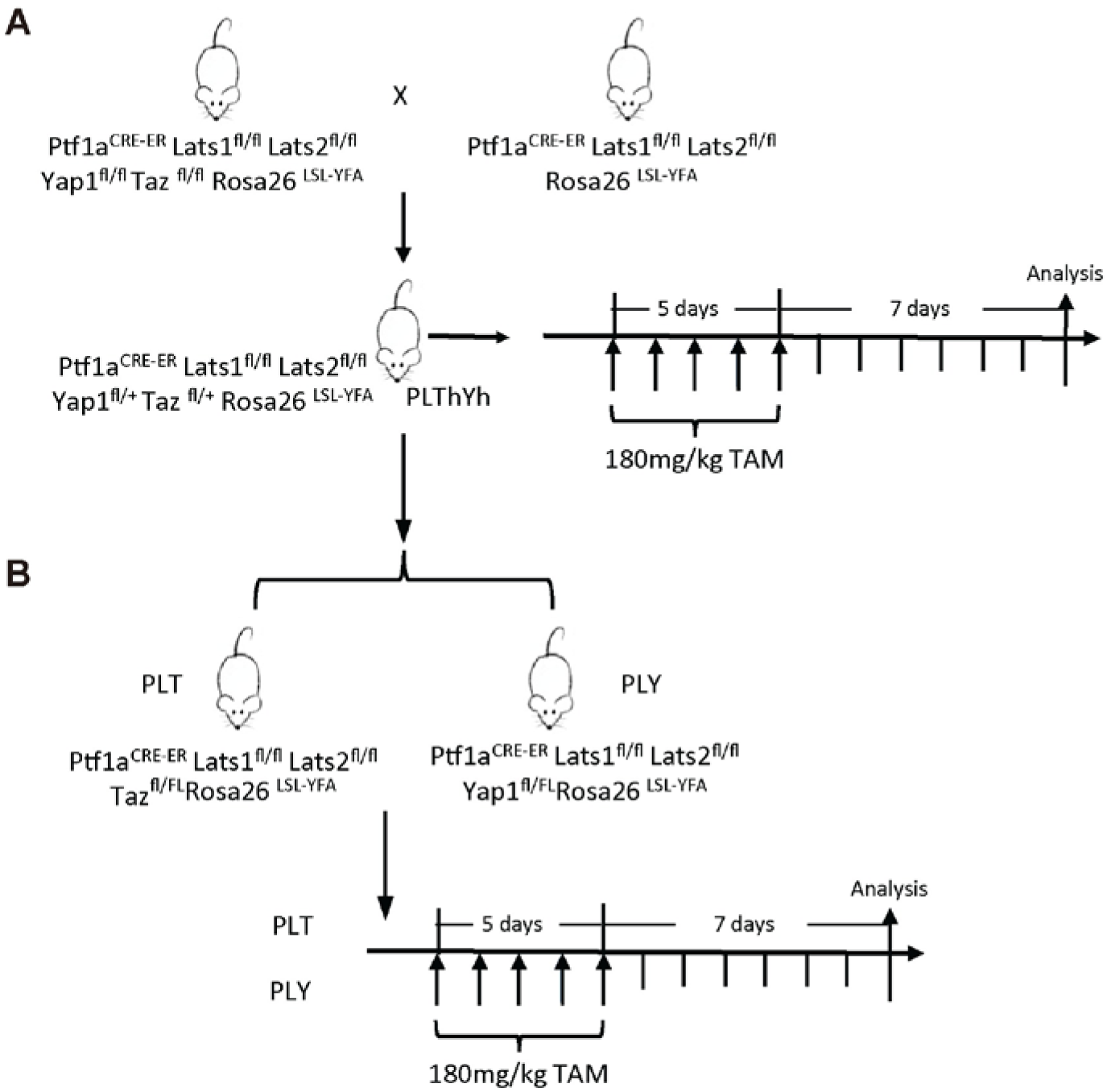
The schema of experimental procedures to generate and analyze PLThYh, PLT and PLY mice.

**Supplementary Figure 2:**
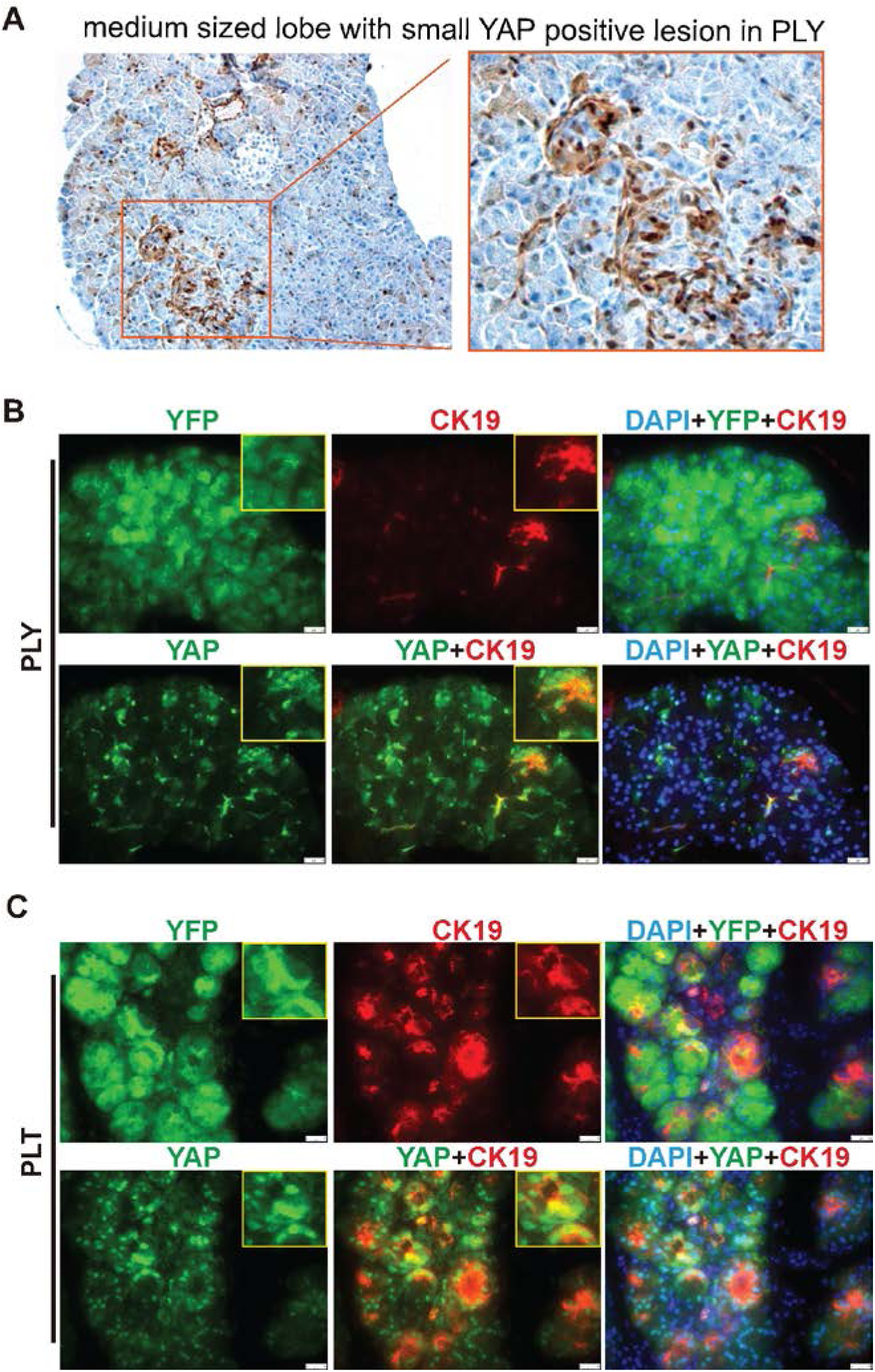
Yap1 expression patterns in ADM-like lesions in PLT and PLY mice. A) The representative IHC staining images of Yap1 in the sporadic lesions in the pancreas collected from PLY mice 7 days after final TAM administration. B) The representative IF images of GFP, Amylase and Yap1 in the pancreas collected from PL and PLY mice 7 days after final TAM administration. C) The representative IF images of GFP, Amylase and Yap1 in the pancreas collected from PL and PLT mice 7 days after final TAM administration.

